# Antibodies to Coagulase of *Staphylococcus aureus* crossreact to Efb and reveal different binding of shared Fibrinogen binding repeats

**DOI:** 10.1101/2022.04.01.486801

**Authors:** Federico Bertoglio, Ya-Ping Ko, Sheila Thomas, Liliana Giordano, Francesca Romana Scommegna, Doris Meier, Saskia Helmsig Polten, Marlies Becker, Srishtee Arora, Michael Hust, Magnus Höök, Livia Visai

## Abstract

*Staphylococcus aureus* pathology is caused by a plethora of virulence factors able to combat multiple host defence mechanisms. Fibrinogen (Fg), a critical component in the host coagulation cascade, plays an important role in the pathogenesis of this bacterium, as it is the target of multiple staphylococcal virulence proteins. Amongst its secreted virulence factors, Coagulase (Coa) and Extracellular fibrinogen-binding protein (Efb) share common Fg binding motives and have been described to form a Fg shield around staphylococcal cells, thereby allowing efficient bacterial spreading, phagocytosis escape and evasion of host immune system responses. Targeting these proteins with monoclonal antibodies thus represents a new therapeutic option against *S. aureus*. To this end, here we report the selection and characterization of fully human, sequence-defined, monoclonal antibodies selected against the C-terminus of Coagulase. Given the functional homology between Coa and Efb, we also investigated if the generated antibodies bound the two virulence factors. Thirteen unique antibodies were isolated from naïve antibodies gene libraries by antibody phage display. As anticipated, most of the selected antibodies showed cross-recognition of these two proteins and among them, four were able to block the interaction between Coa/Efb and Fg. Furthermore, our monoclonal antibodies could interact with the two main Fg binding repeats present at the C-terminus of Coa and distinguish them, suggesting the presence of two functionally different Fg-binding epitopes.

**Importance:** The death toll related to methicillin-resistant *S. aureus* piled to almost 1 million people in only one year (2019), ascribing *S. aureus* to the second leading cause of deaths associated with antimicrobial resistance. Therefore, new therapeutic strategies must be investigated. Blocking the adhesion step with the use of monoclonal antibodies is one promising alternative and Fg is a central plasma protein involved in staphylococcal infection. We present here a panel of monoclonal antibodies raised against Coa, cross-reacting to Efb and targeting the shared Fg binding repeats of Coa. In addition, we describe new epitope determinants in the repeated region of Coa, highlighted by differential binding of the newly selected antibodies.

## Introduction

*Staphylococcus aureus* has a large set of finely-tuned virulence-associated genes that has endowed this bacterium with highly adaptive and versatile strategies to survive in beneficial as well as in hostile environments (1–6). Two major classes of virulence factors belong to <u>C</u>ell <u>W</u>all-<u>A</u>nchored (CWA) adhesins (2) and a group of secreted proteins called <u>S</u>ecretable <u>E</u>xpanded <u>R</u>epertoire <u>A</u>dhesive <u>M</u>olecules (SERAMs) (7). The most represented activity in both groups of virulence factors is their ability to bind fibrinogen (Fg), a host blood glycoprotein. For instance, SERAMs <u>Coa</u>gulase (Coa), <u>v</u>on <u>W</u>illebrand factor <u>b</u>inding <u>p</u>rotein (vWbp), <u>E</u>xtracellular <u>f</u>ibrinogen-<u>b</u>inding protein (Efb), <u>E</u>xtracellular <u>a</u>dhesive <u>p</u>rotein (Eap), <u>E</u>xtracellular <u>m</u>atrix binding <u>p</u>rotein (Emp) all bind Fg (8). Amongst them, prothrombin-activating proteins Coa and vWbp engage Fg independently from prothrombin (9–12). The Fg binding activity of SERAMs, especially well studied for Coa, vWbp and Efb, is mainly located in unordered regions of these proteins (8, 9, 11).

Fg is a large, fibrous plasma glycoprotein with three pairs of polypeptide chains, designated Aα, Bβ and γ. During haemostasis and clot formation, it self-assembles into an insoluble fibrous gel upon conversion to fibrin (13, 14). The role of Fg in bacterial infection has been mainly regarded as protective “haemostatic containment”, owing to the ability of Fg/fibrin to entrap bacteria, reducing their proliferation and dissemination, and fibrin-mediated recruitment of immune cells to clear invading bacteria (15–17). As mentioned so far, *S. aureus* harnesses an impressive array of virulence factors that can interact with Fg. Multiple recent evidence has demonstrated that the interaction with Fg may drive different host responses based on the tissutal context (8). In peritonitis mouse infection models, binding of Fg is fundamental to elicit an antibacterial response and contain infection (18–21). However, the picture is completely reversed in bloodstream infections, where Fg instead promotes spreading of *S. aureus* (22). Therefore, understanding the interactions between *S. aureus* virulence factors and Fg is crucial to understand how new therapeutic opportunities should be designed against the multiple antibiotic-resistant strains of this pathogen.

Efb and Coa are the best characterized SERAM proteins. Efb::Fg interaction is located in the N-terminus half of Efb (23), whereas Coa can bind Fg all throughout its length, with the more potent interactions located in the C-terminus domain (9, 11, 24). Furthermore, both Coa and Efb mediate the formation of a Fg/fibrin shield around staphylococcal cells, thereby protecting bacteria from host immune responses (23, 25–27). Coa also mediates allosteric activation of prothrombin through its N-terminus D1D2 domains promoting fibrin formation (28–30). In respect to therapeutic potential of Coa- and Efb-targeted antibodies, polyclonal rabbit sera raised against Coa (10, 25) or Efb-specific antibodies derived from patients with *S. aureus* infection (31) were able to reduce Fg binding *in vitro* and protect mice from lethal *S. aureus* sepsis.

**Figure 1A** shows the domain organization of Coa protein. The full length protein can be divided into N-terminus and C-terminus halves. N-terminus half of the protein contains D1D2 domains. The C–terminus part of Coa can be divided into two portions: the repeat region of Coa, located at the most C-terminus of the protein, and a linker that connects the D1D2 domain and the repetitive region of Coa. As mentioned earlier, both N-terminus and C-terminus halves can bind Fg but the more potent binding region is located in the C-terminus half. Different recombinant constructs used in the study are also indicated in **Figure 1A**. CoaF contains the linker region and harbours a first slightly divergent and longer repeat, termed CoaR0 (**Figure 1A and C**). The remaining repeats are covered in recombinant construct CoaR, which harbours relatively conserved tandem repeats I-V of 27-residue each. CoaR, together with CoaF, constitutes the C-terminal domain of Coa, expressed as recombinant protein named CoaC. The number of repeats present in Coa protein varies from 1 to 9 copies depending on the *S. aureus* strain, 5 or more being the most common (32). These repeats are shorter than CoaR0, which is the longest repeat able to bind Fg and is present in CoaF, spanning residues from 474 to 505. Therefore, Coa can be divided into several functional domains that have different affinities for Fg (9, 11).

**Figure 1.**
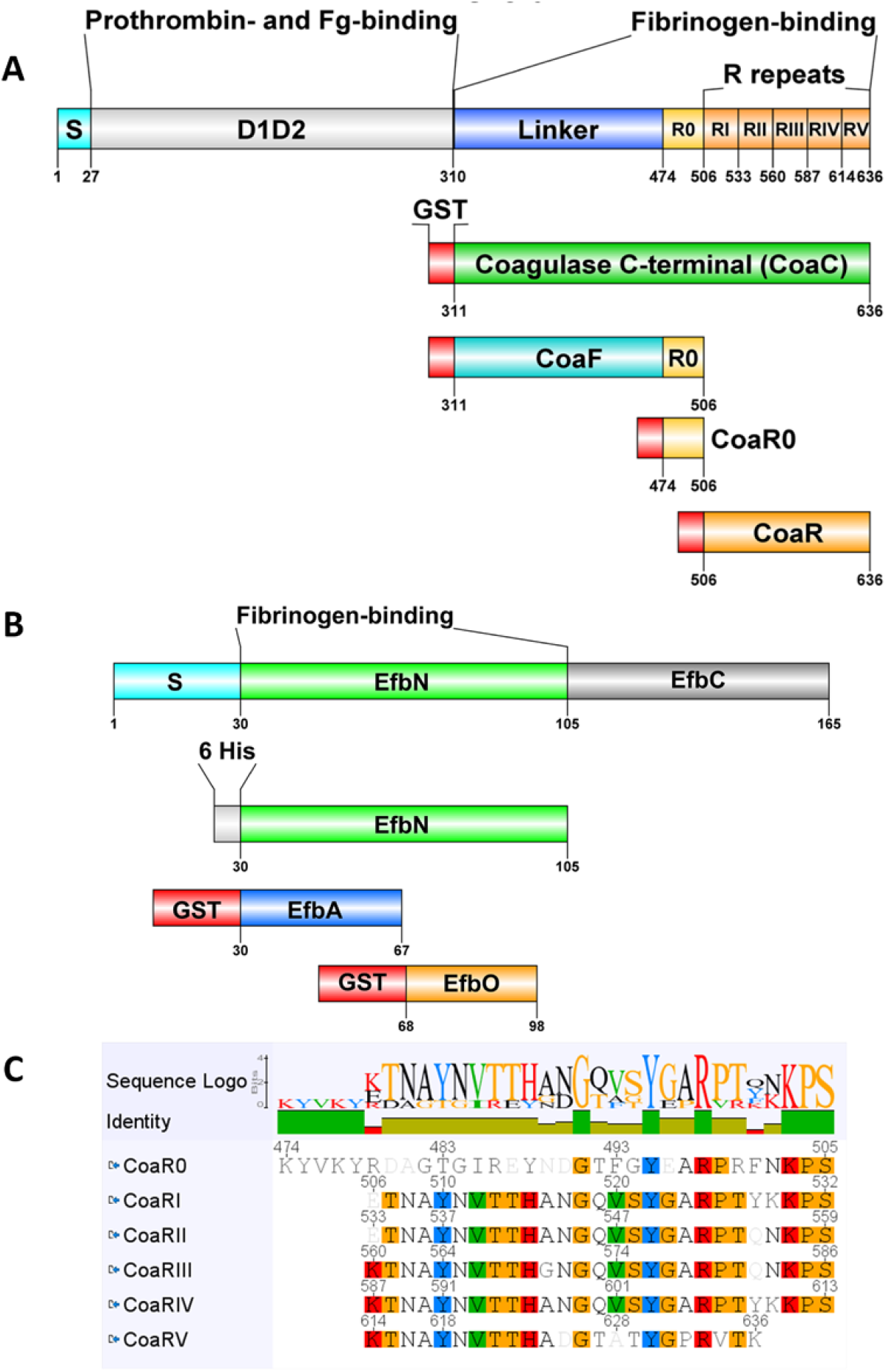
Domain organization and recombinant constructs of Coagulase and Efb. Domains and recombinant constructs of Coagulase (A) and Efb (B) derived from full length protein of *S. aureus* Newman strain are shown. Residues, fibrinogen (Fg)- and prothrombin-binding regions are indicated, Signal peptide (S) is necessary for extracellular release. Gluthation-S-Transferase (GST) tag is in red (not to scale), 6 His tag in grey. Image were prepared with DOG2.0 (65). Panel C shows an alignment, generated with Geneious, of CoaR0 and R repeats of Coa with a sequence logo and identity to highlight the most conserved amino acids. Amino acids in the single repeats are highlighted with Clustal colour scheme if they are present in more than 50% of the sequences.

As mentioned, *S. aureus* Efb also interacts with Fg and belongs to SERAMs (9, 23, 27, 33). It is reported to inhibit complement activation by engaging C3b (34–37), block platelet aggregation (38) and interact with immune cells blocking cellular-mediated immunity (23, 27, 39). Furthermore, Efb can also bind to Complement Receptor 2 on B cells, further tackling adaptive responses of the host (40). Fg-binding activity is located at the N-terminus of Efb and has been mapped to relatively long amino acid stretches termed EfbO and EfbA (**Figure 1B**). The affinity of EfbO for Fg is 200 times higher than that of EfbA, indicating that EfbO is the primary Fg binding site in Efb (23). Coa repeats and Efb N-terminal share homology in their Fg binding mechanisms and likely target the same or overlapping sites in Fg, given that EfbO, CoaR0 and CoaRI peptides are able to inhibit Fg binding of both EfbN and CoaC (9).

The possibility to interfere with Fg binding is thus crucial to understand and block one *S. aureus* pathogenic mechanism. Here, antibodies are not only a tool for blocking Coa and Efb interaction with Fg for research but also potential therapeutic molecules. We used antibody phage display to select several fully human, sequence-defined antibodies against CoaC and characterized them *in vitro*. We found that the anti-CoaC antibodies showed crossreactivity with Efb and were able to discriminate between CoaR0 and CoaRI peptides. In addition, we identified four antibodies that were able to inhibit Coa and Efb Fg-binding.

## Results

### Anti-Coa C antibody selection and production

To select Coa-targeting antibodies, the naïve antibody gene libraries HAL9 (λ repertoire) and HAL10 (κ repertoire) were used as sources for scFv selection by phage display (41). We reasoned that because of the higher affinity of CoaC for Fg than CoaN (9, 11), CoaC would represent a better target to inhibit Fg binding activity (**Figure 1A**). After three panning rounds, monoclonal soluble scFv were expressed from a total of 95 colonies in order to identify specific binders through a screening Enzyme-Linked ImmunoSorbent Assay (ELISA). All clones that gave an Signal-to-Noise ratio > 11 were considered possible binders (**Supplementary Fig. 1**). This selection yielded 45 positive specific hits, termed FBE5 antibodies. No signal was detected both against Bovine Serum Albumin (BSA), used as a negative control, (**Supplementary Fig. 1**) and GST (data not shown). After BstNI digestion, sequencing and analysis with VBASE2 Fab tool (42), 11 unique antibodies were converted in scFv-Fc, an IgG-like divalent format, transiently produced in HEK293.6E cells and Protein A-purified from the clarified supernatant (43). Pure monoclonal <u>A</u>ntibodies (mAbs) preparations were obtained, as indicated by SDS-PAGE (data not shown).

### Anti-Coa C antibody dose-dependent binding to Coa and Efb

The binding of the 11 scFv-Fcs raised against CoaC was further assessed with a titration ELISA, to determine the EC_50._ All FBE5 antibodies bound specifically to CoaC recombinant protein (**Figure 2**) with half-maximum binding in the range between 1,35×10^−8^ M (FBE5-C8) and 5,13×10^−10^ M (FBE5-F11) (**Table 1**).

**Table 1.** Apparent K_d_ for FBE 5 antibodies expressed in M derived from half-maximum binding determined in ELISA. ND: not determinable. NSB: Non-Sigmoidal weak Binding

| ANTIBODY | $K_d$ app (M) for CoaC | $K_d$ app (M) for CoaF | $K_d$ app (M) for CoaR0 | $K_d$ app (M) for EfbN | $K_d$ app (M) for EfbA | $K_d$ app (M) for EfbO |
| --- | --- | --- | --- | --- | --- | --- |
| FBE5-A5 | $5,34 \times 10^{-9}$ | $1,47 \times 10^{-9}$ | $1,86 \times 10^{-9}$ | $3,24 \times 10^{-9}$ | $4,38 \times 10^{-9}$ | $4,27 \times 10^{-9}$ |
| FBE5-A6 | $1,24 \times 10^{-9}$ | $1,15 \times 10^{-9}$ | $1,9 \times 10^{-9}$ | $3,54 \times 10^{-9}$ | $8,54 \times 10^{-9}$ | $1,3 \times 10^{-8}$(NSB) |
| FBE5-A12 | $5,55 \times 10^{-10}$ | $6,47 \times 10^{-10}$ | $2,44 \times 10^{-9}$ | $8,66 \times 10^{-9}$ | $7,28 \times 10^{-8}$(NSB) | $3,51 \times 10^{-8}$(NSB) |
| FBE5-B9 | $1,74 \times 10^{-9}$ | $1,12 \times 10^{-9}$ | $2,19 \times 10^{-9}$ | $3,45 \times 10^{-9}$ | $2,59 \times 10^{-8}$ | $8,27 \times 10^{-9}$ |
| FBE5-C1 | $1,13 \times 10^{-9}$ | $4,59 \times 10^{-9}$ | $2,43 \times 10^{-8}$(NSB) | $2,44 \times 10^{-8}$(NSB) | $4,09 \times 10^{-8}$(NSB) | ND |
| FBE5-C8 | $1,35 \times 10^{-8}$ | $7 \times 10^{-9}$ | $1,93 \times 10^{-7}$(NSB) | $1,7 \times 10^{-7}$(NSB) | ND | ND |
| FBE5-D9 | $7,79 \times 10^{-10}$ | $5,36 \times 10^{-10}$ | $1,14 \times 10^{-9}$ | $1,31 \times 10^{-9}$ | $4,48 \times 10^{-9}$ | $2,68 \times 10^{-9}$ |
| FBE5-D10 | $1,4 \times 10^{-9}$ | $9,5 \times 10^{-10}$ | $8,56 \times 10^{-10}$ | $1,97 \times 10^{-9}$ | $1,57 \times 10^{-9}$ | $3,65 \times 10^{-9}$ |
| FBE5-E5 | $6,27 \times 10^{-9}$ | $9,5 \times 10^{-9}$ | $8,64 \times 10^{-7}$(NSB) | ND | ND | ND |
| FBE5-F9 | $2,85 \times 10^{-9}$ | $1,63 \times 10^{-9}$ | $3,16 \times 10^{-9}$ | $6,01 \times 10^{-9}$ | $2,15 \times 10^{-8}$ | $7,97 \times 10^{-9}$ |
| FBE5-F11 | $5,13 \times 10^{-10}$ | $2,44 \times 10^{-10}$ | $2,13 \times 10^{-10}$ | $2,19 \times 10^{-10}$ | $4,83 \times 10^{-10}$ | $3,96 \times 10^{-10}$ |

**Figure 2.**
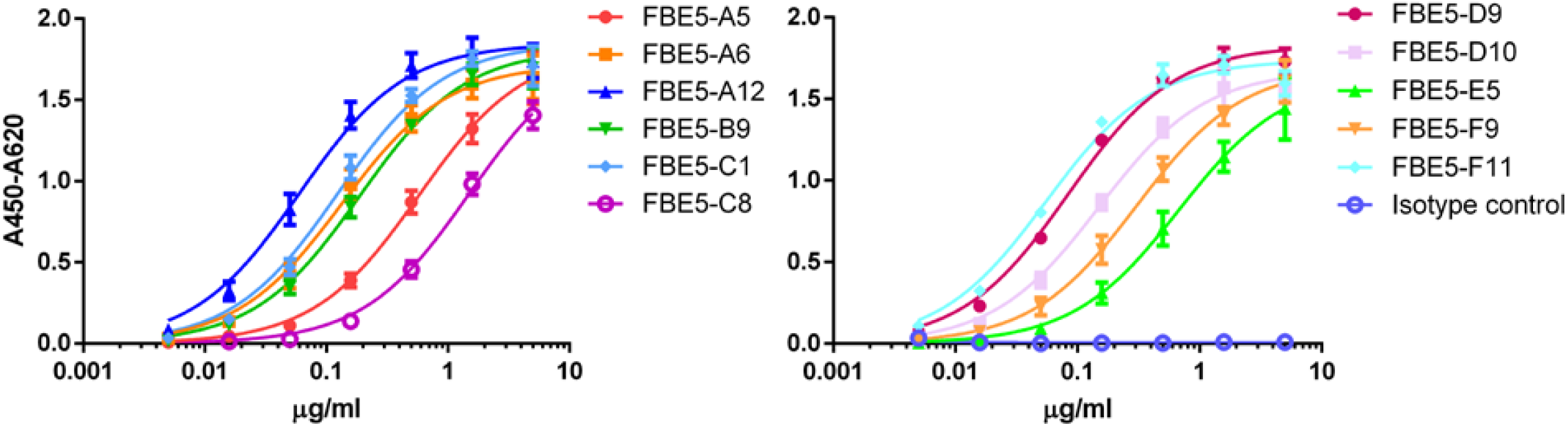
FBE5 antibodies dose-dependently bind CoaC fragment. Titration ELISA to evaluate the binding of FBE5 scFv-Fcs to the antigen CoaC (200ng/well immobilized protein). BSA binding curves were always below 0,1 and are not reported for clarity. Isotype control is an unrelated scFv-Fc with human Fc moiety. Data±SEM are representative of two independent experiments.

Since Coa and Efb share Fg binding motifs, we reasoned that monoclonal antibodies raised against CoaC may crossreact to Efb, specifically to the latter’s N-terminal fragment, where the two functional Fg binding sequences (EfbA and EfbO) are located. Furthermore, CoaC itself harbours a linker region and different Fg binding repeats. Therefore, we wondered if the generated antibodies were able to recognize distinct epitopes within different regions of CoaC and also if any crossreactivity with Efb was detectable. To this end, a single-point ELISA was performed against different recombinant fragments of Coa (namely CoaF, CoaR0, CoaR) (**Figure 1A**) and Efb (EfbN, EfbA and EfbO) (**Figure 1B**). Strikingly, all mAbs bound CoaF and CoaR0 but not CoaR, a construct containing CoaRI-homologous repeats, but not CoaR0 (**Figure 1C and Figure 3**). Similarly, all antibodies, except FBE5-C8 and FBE5-E5, bound to different extents to Efb recombinant constructs tested (**Figure 3**).

**Figure 3.**
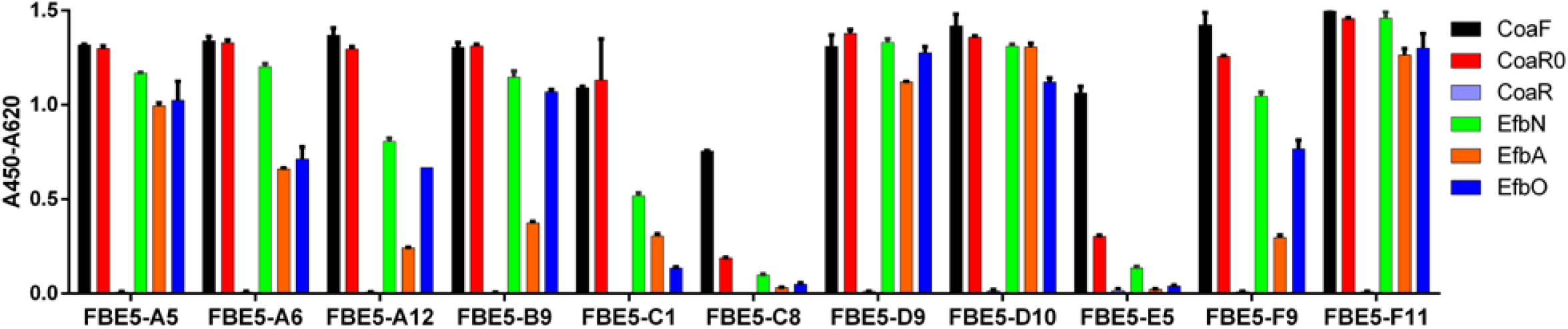
Antibodies generated against CoaC bind CoaR0 repeat, but not CoaRI-RV repeats, and cross react with EfbN. Single point ELISA to evaluate the binding of FBE5 scFv-Fc to fragments of Coa and Efb (200ng/well immobilized protein, [scFv-Fc] = 0,5μg/ml). Represented is the average ±SEM.

To better investigate the binding of each antibody to the different Efb and Coa recombinant proteins, each FBE5 antibody was titrated on CoaF, CoaR0, EfbN, EfbO and EfbA (**Supplementary Figures 2 and 3**) and the respective apparent affinities were calculated (**Table 1**). The best antibody was FBE5-F11, which displayed EC_50_ values in the sub-nanomolar range towards each Coa and Efb construct. The antibodies that showed weak-to-absent binding to all Efb and Coa fragments except CoaF were FBE5-C1, FBE5-C8 and FBE5-E5.

### Generation and characterization of antibodies specific to Coa R

Given that during the previous round of selection none of the characterized antibodies recognized CoaR, another panning was performed specifically to raise antibodies that are able to bind the Coa RI-RV repeats contained in the CoaR fragment (**Figure 1A and 1C**). Isolation of antibodies with this specificity proved particularly ardous in our setting. We screened 380 colonies and were able to retrieve only 10 hits, which upon sequencing revealed to be only two unique antibodies, termed LIG40-A11 and LIG40-D8. Similarly to FBE5 antibodies, the two anti-CoaR mAbs were reformatted in the scFv-Fc divalent format and recombinantly expressed. Dose dependent binding of LIG40 mAbs against Coa and Efb constructs was verified in titration ELISA (**Figure 4**). Both mAbs showed specific high-apparent affinity binding to both CoaR and CoaC proteins, as expected. In particular, LIG40-A11 was specific to CoaR, whereas LIG40-D8 showed binding also to CoaF and to a low level to CoaR0, suggesting a cross-reactivity to CoaR0 repeat. Of note, none of the two antibodies bound to Efb fragments. EC_50_ values against the different Coa constructs for both antibodies are reported in **Table 2** and are almost all in the sub-nanomolar range.

**Table 2.**
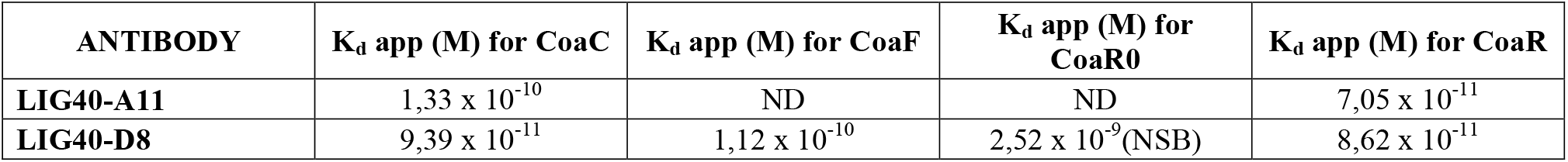
Apparent K_d_ for LIG40 antibodies expressed in M derived from half-maximum binding determined in ELISA. ND: not determinable. NSB: Non-Sigmoidal weak Binding

| ANTIBODY | $K_d$ app (M) for CoaC | $K_d$ app (M) for CoaF | $K_d$ app (M) for CoaR0 | $K_d$ app (M) for CoaR |
| --- | --- | --- | --- | --- |
| LIG40-A11 | $1,33 \times 10^{-10}$ | ND | ND | $7,05 \times 10^{-11}$ |
| LIG40-D8 | $9,39 \times 10^{-11}$ | $1,12 \times 10^{-10}$ | $2,52 \times 10^{-9}$(NSB) | $8,62 \times 10^{-11}$ |

**Figure 4.**
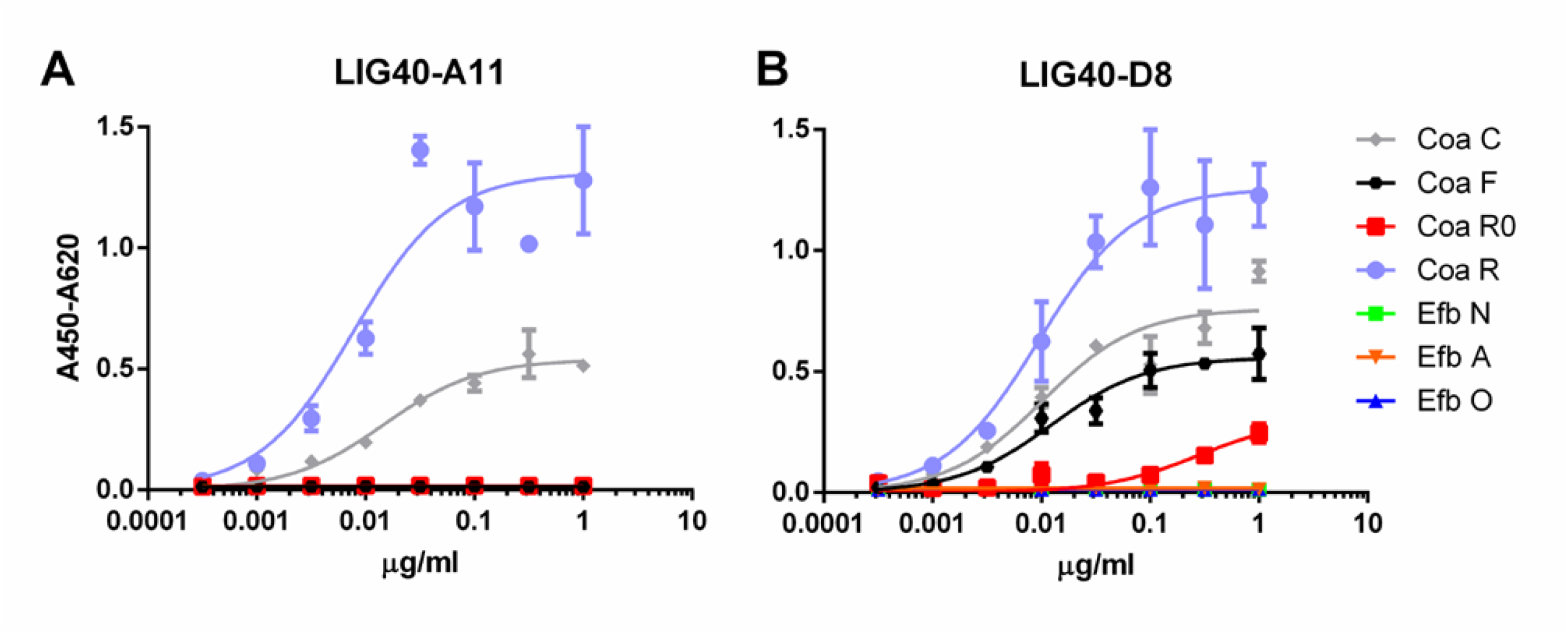
Dose-dependent binding of anti-CoaR mAbs to Coa and Efb recombinant proteins. Titration ELISA to investigate binding of LIG40-A11 (A) and LIG40-D8 (B) to CoaC, CoaR, CoaR0, CoaF, EfbN, EfbA, EfbO recombinant fragments and determine EC_50_. BSA and an unrelated human scFv-Fc (both not represented for clarity) were used as negative and isotype controls, respectively, and showed no binding.

### Inhibition of Coa and Efb fibrinogen binding by the selected mAbs

Since we showed that FBE5 and LIG40 mAbs bind functional Fg-binding Coa fragments and FBE5 mAbs also bind Efb fragments, we investigated if these mAbs can block the interaction between Fg and their antigens. Binding of CoaF, CoaR0, EfbN, EfbA and EfbO to purified human Fg, which was immobilized on an ELISA plate, was assessed in the presence of increasing concentrations of FBE5 antibodies. CoaR was not tested since none of FBE5 mAbs did recognize CoaR. FBE5-A12, FBE5-D10, FBE5-F9 and FBE5-F11 showed the best dose-dependent inhibition of Fg binding in good accordance with binding data (**Figure 5**). Specifically, FBE5-F11 antibody was the most potent inhibitor of Fg binding to all Coa and Efb recombinant proteins tested, reaching an almost complete inhibition of CoaF binding to Fg at 5 μg/ml. Similarly, below 20% of Coa R0 residual binding to Fg was detected at 5 μg/ml of FBE5-F11. The same antibody inhibited EfbN, EfbA and EfbO binding to Fg less efficiently, resulting in more than 60% inhibition only at the highest concentration tested. FBE5-A12, FBE5-D10 and FBE5-F9 inhibited binding of Coa fragments to Fg more than binding of Efb. In particular, FBE5-A12 showed an inhibition of CoaF comparable to FBE5-F11, but was less effective against CoaR0. FBE5-D10 and FBE5-F9 showed inhibition only at high concentration (50 μg/ml) of both CoaF (more than 70% and almost 100%, respectively) and CoaR0 (more than 70% for both mAbs). FBE5-A12, FBE5-D10 and FBE5-F9 displayed a dose-dependent inhibition of only EfbA construct, with no remarkable inhibition of EfbN and EfbO proteins. It is however to be highlighted that EfbA harbours a less potent Fg binding sequence (23).

**Figure 5.**
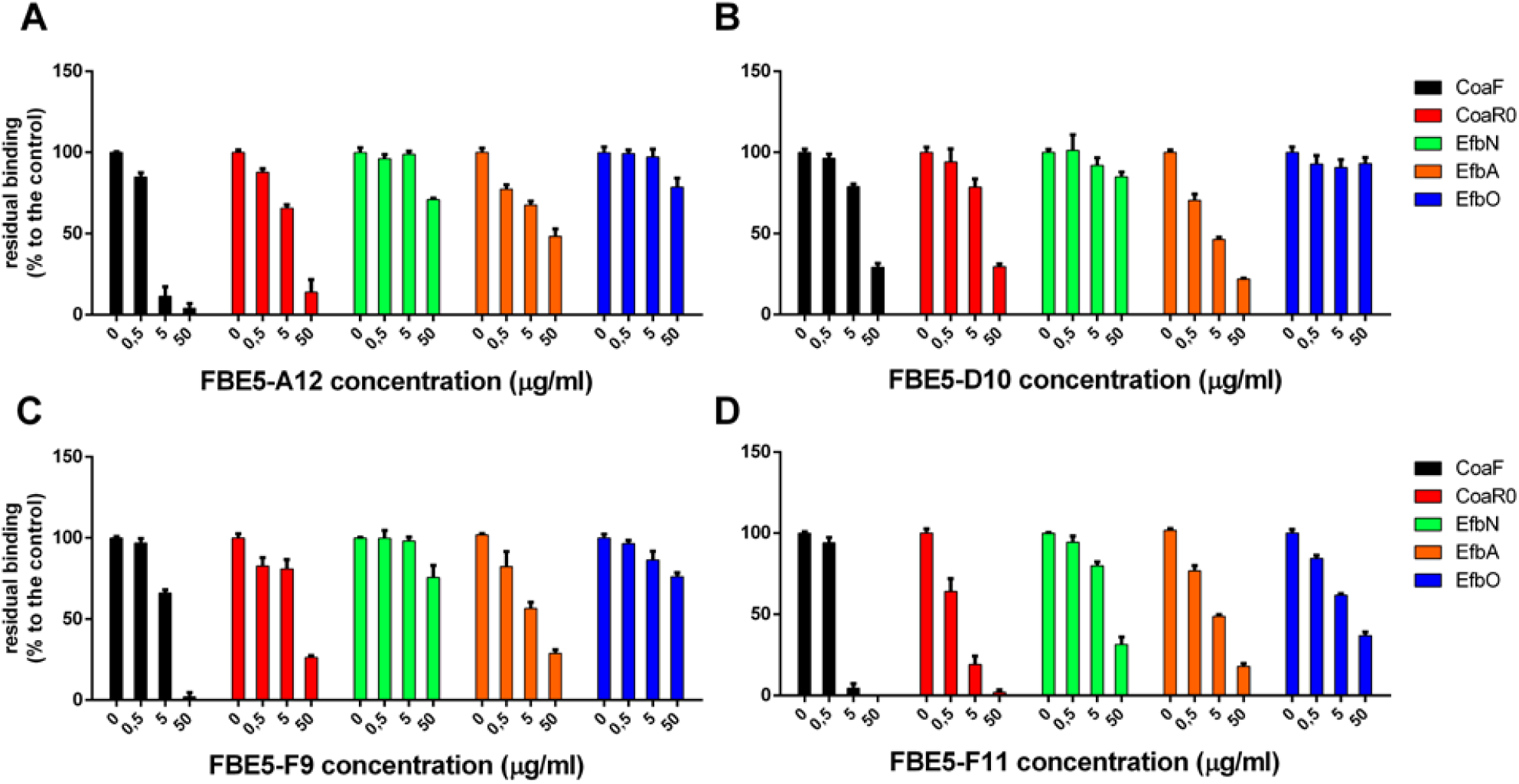
Anti-CoaC antibodies inhibit both Coa and Efb fibrinogen binding activity. Antibodies FBE5-A12 (A), FBE5-D10 (B), FBE5-F9 (C) and FBE5-F11 (D) were pre-incubated at the indicated amounts with Coa or Efb recombinant constructs (CoaF, CoaR0, EfbN and EfbO at final concentration of 10nM, EfbA at 750μM) and then transferred to a Fg-coated ELISA plate. The remaining Fg-bound antigens were detected through their tags (GST, except for EfbN which harbours a 6xHis tag). Control wells with no antibody were set to 100% and the residual binding of Coa and Efb constructs was determined by comparing control wells with the ones where indicated amounts of mAbs were added. Average ± SEM of two independent experiments is represented.

The remaining 7 antibodies showed limited-to-no inhibition of Coa fragments at high concentration and essentially displayed no inhibition against Efb protein (**Supplementary Fig. 4**). Similarly, both LIG40 antibodies were tested for inhibition of Fg binding to CoaC, CoaF, CoaR0 and CoaR but did not show any inhibiting activity (**Supplementary Fig. 5**).

### Binding of mAbs is affected differently by CoaR0 and CoaRI peptides

In order to better understand if the generated monoclonal antibodies engage at their epitope the 2 Fg binding motives of Coa, peptides corresponding to CoaR0 and CoaRI repeats (**Figure 1C**) of *S. aureus* strain Newman were synthetically manufactured and used to challenge binding of both FBE5 and LIG40 mAbs to their respective antigens. To this end, a competition ELISA was performed to evaluate the binding of a fixed concentration of antibody to immobilized Coa constructs in the presence of increasing concentrations of either peptide CoaR0 or CoaRI. The fixed quantity of antibody chosen allowed to detect sufficient binding of antibodies, yet to be able to see any variation upon addition of the peptides. Peptide CoaR0 inhibited FBE5 antibodies binding to CoaC in a dose-dependent manner (**Figure 6, panels A** and **B**) whereas CoaRI had no effect (**Figure 6, panels C** and **D**). This result corroborates the binding data that showed recognition of CoaF and CoaR0 fragments, but not of CoaR (**Figure 3** and **Supplementary Fig. 2** and **3**). CoaF and CoaR0 do indeed contain CoaR0 repeat, which is conversely absent in CoaR, where repeats similar to CoaRI peptide are located. These data suggest that all FBE5 antibodies bind epitopes within CoaR0.

**Figure 6.**
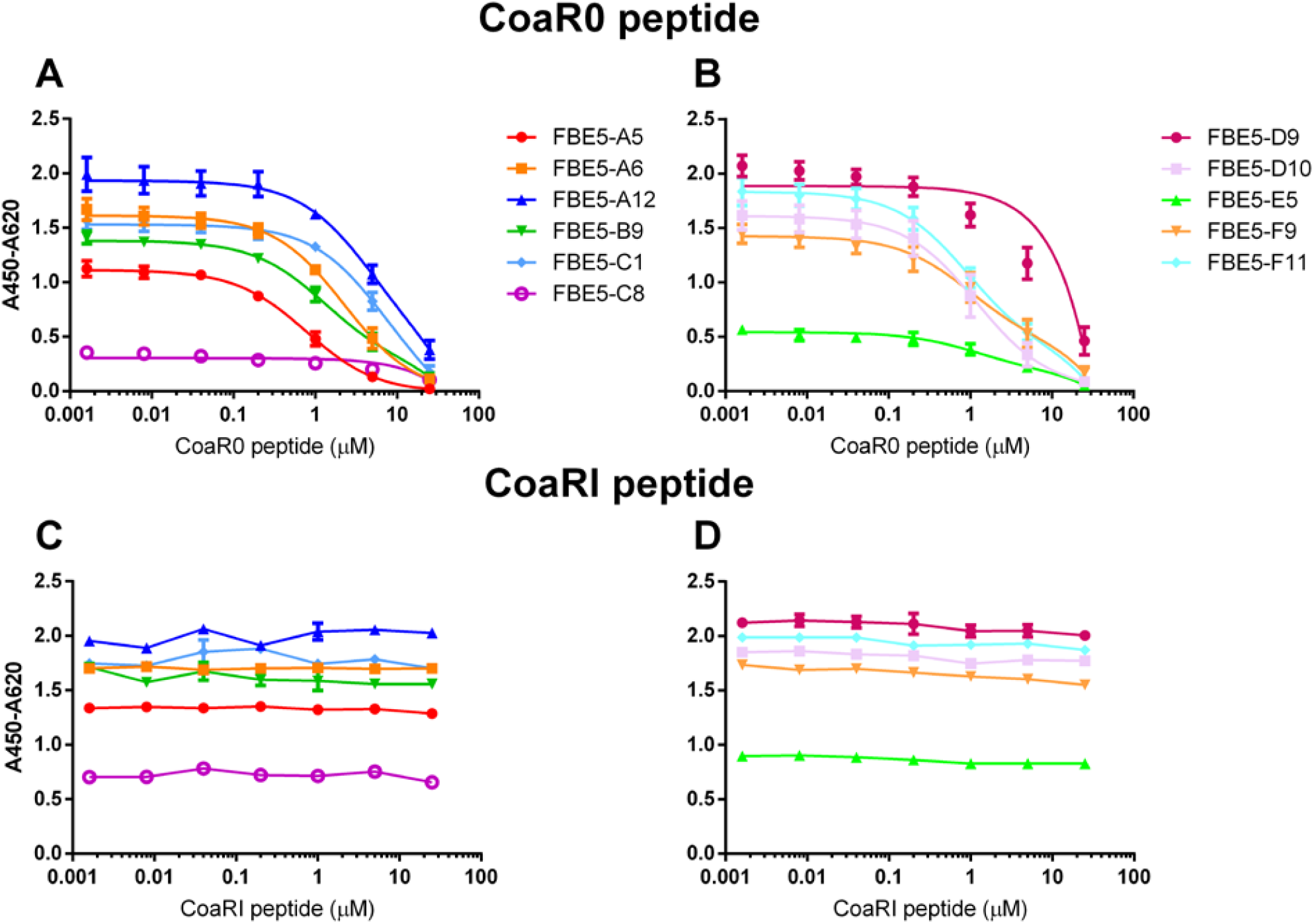
Binding of FBE 5 mAbs to Coa C is inhibited by CoaR0 peptide but not by CoaRI peptide. CoaC was immobilized on an ELISA plate and a fixed quantity of each mAb was added (0.5μg/ml) with different dilutions of either CoaR0 (A and B) or CoaRI (C and D) peptide. Detection of mAb was performed through anti-human HRP (HorseRadish Peroxidase)-conjugated secondary antibody. Data ± SEM are reported and are representative of two independent experiments.

The effect of CoaR0 and CoaRI peptides was investigated also on CoaC and CoaR binding of both LIG40 mAbs. Surprisingly, LIG40-A11 and LIG40-D8 behaved differently in the presence of the two peptides (**Figure 7**). First and most importantly, LIG40-A11 was inhibited only by CoaRI peptide, when tested against both CoaC and CoaR proteins (**Figure 7, panels A** and **C**). In a symmetrical opposite way, LIG40-D8 was only impaired in its binding activity by CoaR0 peptide (**Figure 7, panels B** and **D**). Secondly, to achieve appreciable inhibition, high concentration of peptides needed to be used for both antibodies (above 10 μM). These results show that LIG40-D8 targets an epitope similar to CoaR0 peptide, yet present in CoaR, which harbours only CoaRI-type repeats. On the other hand, LIG40-A11 binds to an epitope specific of CoaR repeats.

**Figure 7.**
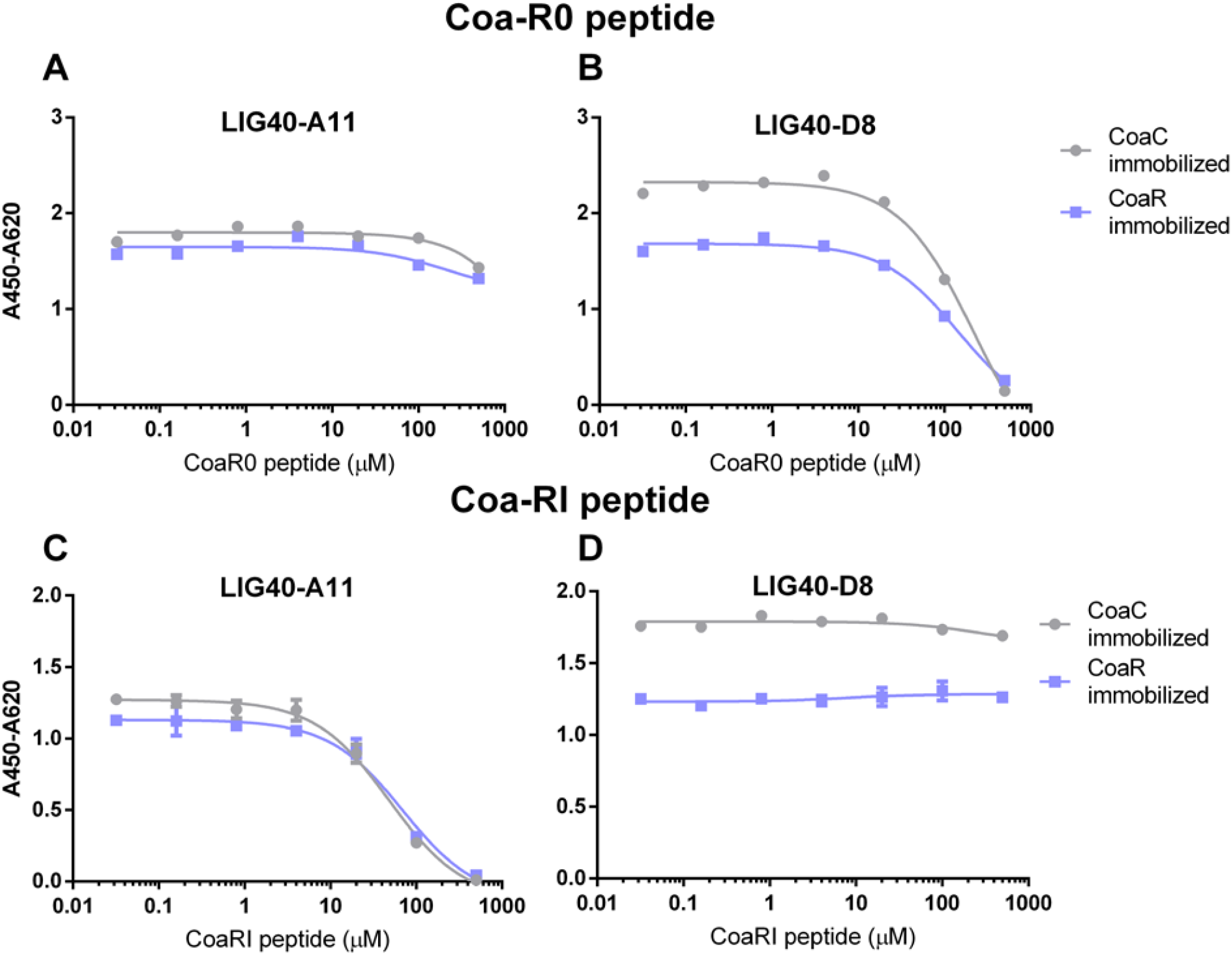
Anti-CoaR antibodies are inhibited differently by CoaR0 and CoaRI peptides. CoaC and CoaR were immobilized on an ELISA plate. 0.5ug/ml of LIG40-A11 (A and C) or LIG40-D8 (B and D) were incubated with different dilutions of either CoaR0 (A and B) or CoaRI (C and D) peptides. Detection of mAb was performed through anti-human HRP-conjugated secondary antibody. Data ± SEM are reported and are representative of two independent experiments.

Collectively, these data show that fully human, sequence-defined, monoclonal antibodies against Coa C-terminal fragment were able to engage and block Fg-binding motives both in Coa and Efb. Furthermore, we showed that it is possible to discriminate between CoaR0 and CoaRI repeats through monoclonal antibodies.

## Discussion

*S. aureus* has for a long time been a critical global healthcare threat owing to increase in spread and virulence of antibiotic-resistant strains (44, 45). The discovery and introduction of radically new antibiotic classes into the market has been lagging for two decades (46). Therefore, new approaches to tackle *S. aureus* infections are of foremost interest. A first crucial aspect in *S. aureus* pathogenesis is the attachment to host tissues. Among these interactions, Fg seems to play a dominant role (8). Indeed, *S. aureus* has evolved a vast arsenal of proteins to interact with this soluble plasma protein: first, <u>M</u>icrobial <u>S</u>urface <u>C</u>omponents <u>R</u>ecognizing <u>A</u>dhesive <u>M</u>atrix <u>M</u>olecules (MSCRAMMs) are cell wall-bound proteins primarily involved in extracellular matrix components binding to secure bacterial adhesion to host tissues (2). A second class is collectively referred to as SERAMs proteins, which are secreted and still interact with several extracellular matrix molecules displaying also an immune evasion and dissemination function (7). Among MSCRAMMs, ClfA and ClfB, FnbpA and FnbpB and SdrE/Bbp bind different segments of Fg molecule. Fg-binding activity is also prominent in SERAMs, where unordered regions of Coa, vWbp, Efb, Eap and Emp present Fg binding as a common feature. The comprehensive picture of these proteins seems not yet fully disclosed, as the recent initial characterization of vhp shows (47).

Also the role of Fg is at the crossroad between its well described role in haemostasis and its importance in mediating immune response (14, 48). Much preclinical evidence also showed that mutated versions of Fg cannot efficiently clear infection mediated by *S. aureus* thus compromising immune response towards the pathogen (19–21, 49). Furthermore, preclinical studies together with vaccine candidates have shown that ClfA-mediated Fg interaction is a viable alternative for possible therapeutic strategies (50–52). Therefore, all these presented interactions show how intricate the interaction with *S. aureus* and Fg is and thus its extremely high potential as a therapeutic target for alternative treatment strategies.

To this end, anti-Coa antibodies have been generated by antibody phage display, using human naïve libraries (41), providing sequence-defined mAbs: 11 mAbs with unique sequences recognized CoaF and CoaR0 fragments upon panning against CoaC (FBE5 mAbs) and 2 mAbs directed against CoaR (LIG40 mAbs). Of note, to obtain the latter antibodies, 4 times our usual number of clones had to be screened, to obtain only 10 hits and in the end 2 unique clones. In comparison, selection of FBE 5 mAbs had a higher hit rate (45 positive hits/95 colonies screened). This may be a consequence of the unstructured organization of CoaR (11, 30), even if selection was performed on ELISA plates which should “immobilize” antigens in a fixed position. Further support of the unordered nature of the Fg-binding portion of Coa and Efb is given by AlphFold databases (Uniprot entries P07767 and P0C6P2) (53, 54). Interestingly, these structural predictions show that the Fg-binding sequences appear to be more folded than the completely unordered domain surrounding them, in apparent contradiction with prior experimental evidence (11, 30). While AlphaFold predictions pave the way to new questions on how Fg is exactly bound by Coa and Efb, experimental validation of *in silico* data is necessary to draw any solid conclusion.

Since Coa shares sequence and functional homology to Efb, cross-recognition of the generated antibodies was investigated. All FBE5 antibodies showed binding to both Coa and Efb fragments (**Figure 3** and **Supplementary Figures 2** and **3**). In particular, all FBE5 mAbs bound to different extents CoaF, CoaR0, EfbN, EfbA and EfbO fragments, except FBE5-C8 and FBE5-E5 that showed low-affinity binding to Coa and substantially no binding to Efb. Among them, FBE5-F11 had the highest affinity (**Table 1**) and showed the greatest inhibitory effect on Fg binding to CoaF, CoaR0, EfbN, EfbA and EfbO constructs (**Figure 5**). FBE5-A12, FBE5-D10 and FBE5-F9 could also efficiently inhibit all EFb and Coa fragments tested, although to a lower extent than FBE5-F11 (**Figure 5**). This activity correlated with their apparent affinities determined in ELISA (**Table 1**). FBE5-A5, FBE5-A6, FBE5-B9, FBE5-C1 and FBE5-D9, instead, showed inhibition of CoaF, CoaR0 and EfbA (**Supplementary Figure 4**). Essentially, these antibodies were able to inhibit CoaR0-mediated Fg binding, since EfbA is unlikely to be the most functionally relevant Fg-binding region in Efb, given its low affinity for Fg (1μM) (23). Finally, FBE5-C1, FBE5-C8 and FBE5-E5 displayed only minor blocking activity on EfbA, matching the apparent affinity measurement of these antibodies.

Concerning LIG40 antibodies raised against CoaR, they did not show any cross-reaction to Efb. Surprisingly, both of them could not inhibit Fg-binding of both CoaC and CoaR (**Supplementary Figure 5**), despite their high affinity binding to functional Fg-binding sequences CoaR0 and CoaRI (**Figure 4** and **Table 2**). When binding of LIG40-A11 and LIG40-D8 to CoaC and CoaR was challenged with synthetic CoaR0 and CoaRI peptides (**Figure 7**), high concentrations of peptides were necessary for competition. This could indicate, on one side, that the epitope is not properly represented in the peptide. On the other hand, it is very plausible that given the repetitive nature of CoaR, multiple binding sites for these antibodies are available within the same construct, thus higher concentration of peptide is needed in order to exert an competitive effect. It is also highly unlikely that a single mAb’s paratope could span the entire linear 27 amino acid-long Fg binding repeat. These considerations hint that the inability of these antibodies to block Fg binding might be due to the repetitive nature of CoaR.

Furthermore, CoaR0 peptide could inhibit all FBE5 and, surprisingly, LIG40-D8 mAbs binding to CoaC, instead had no effect on LIG40-A11 binding to both CoaR and CoaC. Conversely, CoaRI peptide did inhibit LIG40-A11 binding, leaving unaffected the binding of all FBE5 and LIG40-D8 mAbs. The latter antibody showed indeed binding, albeit weaker, to CoaF and CoaR0 constructs (**Figure 4, Table 2**), even though its selection was performed on CoaR, which does not contain CoaR0 repeat. These results together suggest that these antibodies are targeting different epitopes in CoaR and that Fg binding repeats may assume similar conformations, however representing two functionally distinct epitopes in Coa. It could be speculated that LIG40-D8 is targeting conserved residues present both in CoaR0 and CoaRs repeats. Their role needs further clarification since it is clear that the binding site of CoaR0 and CoaRI in the Fg molecule is similar or overlapping. Both peptides are indeed able to inhibit Coa binding to Fg (9). Recent evidence has validated these previous results, highlighting the role of the CoaR0 repeat in Fg binding, further confirming that both CoaR0 and CoaRI are indeed the functional Fg binding repeats. It was also shown that increasing the number of Fg binding repeats does not lead to a cooperative effect and the stoichiometry remains 1:1 (number of repeats: Fg D molecule) (55). This is in further support of the possibility that more than one antibody molecule is necessary to efficiently inhibit Fg binding by all CoaRI-similar repeats, thereby giving a possible explanation why no efficient inhibition could be seen by antibodies directed to CoaR.

Other antibodies that bind either Coa or Efb have been reported. Thomer and colleagues (24) generated 13 mouse monoclonal antibodies by hybridoma technology targeting Coa and investigated two of them (5D5 and 3B3) *in vivo* in a mouse model of *S. aureus* bacteraemia, with no detailed biochemical characterization. 5D5 mAb was assessed to bind the D1 domain of Coa and 3B3 bound the domain containing R repeats. No analysis of their crossreaction with Efb was provided. The only information available about crossreactivity is that no binding to vWbp and IsdA was detected. MAb 3B3 proved its clear efficacy in the bacteraemia mouse model, further highlighting clinical relevance of the repeated region of Coa (24). A detailed biochemical analysis of these antibodies would provide orthogonal confirmation to our results, also in respect to the hypothesis of two classes of motives by CoaR0 and CoaRI repeats. It is also a possibility that the efficacy of mAb 3B3 could be due to the parallel targeting of Coa and Efb. A clear obstacle for 5D5 and 3B3 clinical translation is their fully murine origin.

Shannon and colleagues found that antibodies against Efb from patient sera could be neutralizing *in vitro* and also crossreacting to Coa (56). A further peculiar class of antibodies against Efb, named catalytic antibodies, were isolated (57). This discovery led to the hypothesis that Efb could also act as a B cell superantigen. Another group instead focused on the selection and characterization of recombinant divalent (Fab)2 mAbs from a synthetic phage display library against Efb C-terminal domain (58). The latter work showed both the presence of antibodies specific to Efb C-terminal in patient sera and also that blocking Efb interaction with C3b with the selected divalent mAbs improved mice survival in an infection model.

This present work and research from other groups shows how pivotal may be blocking the multiple activities of proteins engaging Fg, further strengthening a possible therapeutic strategy involving Coa and Efb (59). To the best of our knowledge, this is the only report that investigates these two proteins as potential targets for generation of monoclonal antibodies. None has provided sequenced-defined human mAbs to the Fg-binding domain of Efb and Coa. The use of combination(s) of antibodies directed against either N- or C-terminal of both these proteins most presumably will result in additive effect in inhibiting *S. aureus* pathology. Selection of such antibodies is already underway.

## Material and Methods

### Recombinant proteins and Fg

CoaC, CoaR, CoaF, CoaR0, EfbA and EfbO harbour an N-terminal GST tag, whereas EfbN has been expressed with a 6 His N-terminal tag and the respective expression and purification protocols were previously reported (9, 23). Human Fg was purchased from Enzyme Research and further purified by size exclusion chromatography to eliminate contaminating fibronectin.

### Selection of scFv antibody fragments (panning)

The selection was performed in ELISA plates (Costar), as described earlier (60). In short, 1μg/well of CoaC or CoaR for each of the three panning rounds was immobilized. The immobilisation conditions in this whole work were at 4°C overnight in 50 mM sodium carbonate, pH 9.6. After blocking with 2%(w/v) Milk powder (M) dissolved in PBS 1x + 0.05% Tween20 (PBST), 5 × 10^10^ phage particles from each of both HAL9 and HAL10 hyperphage-packaged naïve antibody gene libraries were used (41, 61). After incubation in the antigen-coated well, stringent washing with PBST was performed by an ELISA washer (Tecan). Phages were eluted with trypsin (10μg/ml in PBS).

*E. coli* TG1 (Lucigen) in 2xYT medium (yeast extract 1% w/v, tryptone 1.6% w/v, NaCl 0.5% w/v) at OD_600_ of 0.5 were infected with eluted phages for 30 min at 37°C and subsequent 30 minutes at 37°C, 500rpm. Cultures were pelleted, resuspended in 2xYT-GA (2xYT with 100mM glucose and 100μg/ml ampicillin) and, upon OD_600_ of 0.4-0.6, infected with M13K07 helper phage (62). Phage particles were produced at 30°C and 500 rpm overnight in 2xYT with 100μg/ml ampicillin and 70μg/ml kanamycin. After centrifugation, the phage-containing supernatant was used for the next panning round. After the third panning round, instead, *E. coli* XL1Blue MRF’ (Stratagene) at OD_600_ of 0.5 in 2xYT with 20 μg/ml tetracycline was infected with eluted phages, plated on 2xYT-AG agar and cultivated overnight at 37°C.

### Production of soluble scFv in microtiter plates

95 or more colonies per each panning were picked and the corresponding 96-well masterplate inoculated in 2xYT-AG and grown overnight at 37°C and 250 rpm. A subculture in 2xYT-AG was incubated at 37°C, 250 rpm for 90 minutes. Cells were pelleted and resuspended in 2xYT with 100μg/ml ampicillin and 50 μM IPTG and cultured at 30°C, 250 rpm overnight.

### Screening ELISA for monoclonal binder identification

High-binding ELISA plates were coated with 2 μg/ml solution of CoaC or CoaR. As negative controls, BSA and GST were tested. The coated plates were blocked with 2%MPBST and washed with MilliQ + 0.05% Tween 20 in an ELISA washer (BioTek). Crude supernatant containing scFv diluted 1:1 with 2%MPBST was transferred to the corresponding well of both the antigen-coated and the control plates. As primary antibody, α-c Myc tag (9E10, in–house production) was diluted 1:1000. The primary antibody was detected with α-mouse IgG HRP (HorseRadish Peroxidase)-conjugated antibody (A0168, Sigma), diluted 1:50000 in 2%MPBST. Development was performed through the substrate Tetramethylbenzidine (TMB). The reaction was stopped adding 0.5M H_2_SO_4_. The plates were read in an ELISA reader (Tecan) at 450 nm and as a reference wavelength 620 nm. The represented data (A450-A620) are the subtraction of the Absorbance (A) at 450 nm (A450) minus those at 620 nm (A620).

### Colony PCR and BstNI digestion of the PCR product

The scFv gene of positive hits was amplified with primers MHLacZ-Pro_f (5’- GGCTCGTATGTTGTGTGG-3’) and MHgIII_r (5’- CTAAAGTTTTGTCGTCTTTCC-3’). The PCR products were analyzed through capillary gel electrophoresis with the QIAxel instrument (Qiagen). The cPCR-amplified scFv gene was then digested with BstNI endonuclease to obtain and compare the band patterning of each scFv amplified gene. Digestion products were analyzed with the QIAxel (Qiagen). Unique binders were then confirmed by Sanger sequencing and VBASE database analysis (42).

### Cloning of scFv gene into vector pCSE2.6-hIgG1-Fc-XP for scFv-Fc expression

The scFv gene was digested from pHAL30 phagemid with NcoI-HF™ and NotI-HF™ (New England BioLabs), separated by agarose gel electrophoresis and DNA was recovered with QIAquick Gel Extraction Kit (Qiagen), according to supplier instructions. The scFv gene was then ligated into pCSE2.6-hIgG1-Fc-XP vector (43) using T4 Ligase (Promega) and transformed into *E. coli* XL1Blue MRF’, according to standard procedures (63). Correct insertion was confirmed by Sanger DNA sequencing, using softwares FinchTV (Geospiza, Inc.) and Multalin (64).

### Mammalian cell transfection, transient expression and purification of scFv-Fc fusions

ScFv-Fcs were produced as described (43) with minor modifications. In particular, purification was performed with a vacuum manifold (Macherey-Nagel) and a 24 deepwell filter plate loaded with MabSelect SuRe™ (rProtein A, GE Healthcare Life Sciences), according to manufacturer instructions. Buffer exchange to PBS was performed with Zeba™ Spin Desalting columns (Thermo Scientific). Protein purity was checked through SDS-PAGE, using standard protocols (63).

### ELISA assays

High-binding ELISA plates were coated with 200 ng/well of indicated recombinant proteins (CoaF, CoaR0, CoaR, EfbN, EfbA and EfbO or BSA for negative controls). After blocking with 2%BSA in PBST and washing with PBST, scFv-Fc in 2%BSA-PBST were incubated on the immobilized proteins. ScFv-Fc were revealed thanks to a polyclonal α-human IgG HRP-conjugated Ab (P0214, Dako), diluted 1:10000. Final development was performed through SigmaFAST-OPD tablets (P9187, Sigma), following producer instructions. Absorbance was recorded in a microplate reader (Clariostar®, BMG-Labtech). Apparent Kd values were obtained through analysis of half maximum binding using GraphPad Prism 6 software (non-linear regression fit).

For inhibition ELISA, 0,25μg/well of Fg were immobilized. Indicated amounts of scFv-Fcs were pre-incubated in a separate plate with a constant concentration of Coa or Efb fragments. Specifically, CoaF, CoaR0, EfbN and EfbO were at a fixed final concentration of 10nM, whereas EfbA was at 750μM. The pre-incubated mixture of Coa/Efb and anti-Coa scFv-Fc was transferred onto the BSA-blocked Fg-coated plate. After incubation and washing, residual bound Coa and Efb were detected with anti-tag HRP-conjugated antibodies diluted 1:10000 in 2%BSA-PBST: α-HIS-tag antibody (A7058, Sigma) for EfbN; α-GST-tag antibody (600-103-200, Rockland) for all other constructs. Development and acquisition were performed as indicated above. Binding of Coa and Efb fragments to Fg (no mAb control) was set to 100% and residual binding to Fg of Coa and Efb fragments in the presence of different concentrations of antibodies was calculated and represented.

For competition ELISAs with CoaR0 and CoaRI peptides, indicated constructs (200 ng/well) were immobilized. Fixed concentration of mAbs (0,5 μg/ml) was added to the wells with indicated amounts of CoaR0 and CoaRI peptides. Detection of residual mAbs bound was performed as mentioned above.

### Peptides

CoaR0 and CoaRI peptides were purchased from Shanghai Hanhong Scientific Co., Ltd. All the peptides were purified using high-performance liquid chromatography and were >95% pure.

## Acknowledgments

FB acknowledges a research travel grant from SIB (Società Italiana di Biochimica e Biologia Molecolare - Italian Sociaety of Biochemistry and Molecular Biology-) useful in the final stages of this study. This research was supported by a grant of the Italian Ministry of Education, University and Research (MIUR) to the Department of Molecular Medicine of the University of Pavia under the initiative “Dipartimenti di Eccellenza (2018–2022)”. The funders had no role in study design, data collection and interpretation, or the decision to submit the work for publication.

## Supplementary Figures Legends

**Supplementary Fig. 1- Screening ELISA results after 3 panning rounds on CoaC**

Both signals (Optical Density 450-620nm) on CoaC and BSA for each of the 95 tested clones are represented as sided bars.

**Supplementary Fig. 2- Dose-dependent binding of anti-CoaC mAbs to Coa recombinant proteins**

Titration ELISA to investigate binding of FBE5 mAbs to CoaF (A) and CoaR0 (B) recombinant constructs and determine EC50. BSA (not represented) and an unrelated human scFv-Fc were used as negative and isotype controls, respectively.

**Supplementary Fig. 3 Dose-dependent binding of anti-CoaC mAbs to Efb recombinant proteins**

Titration ELISA to investigate binding of FBE5 mAbs EfbN (A), EfbA (B), EfbO (C) recombinant constructs and determine EC50. BSA (not represented) and an unrelated human scFv-Fc were used as negative and isotype controls, respectively.

**Supplementary Fig. 4 - Anti-CoaC antibodies not inhibiting both Coa and Efb fibrinogen binding activity**

Antibodies FBE5-A5 (A), FBE5-A6 (B), FBE5-B9 (C) and FBE5-C1 (D), FBE5-D9 (E), FBE5-C8 (F), FBE5-E5 (G) were pre-incubated at the indicated amounts with the fixed amounts of Coa or Efb recombinant constructs (CoaF, CoaR0, EfbN and EfbO at final concentration of 10nM, EfbA at 750μM) and then transferred on a Fg-coated ELISA plate. The remaining Fg-bound antigens were detected through their tags (GST, except for EfbN that harbours a 6xHis tag). Control wells in which no antibody was added were set to 100% and the residual binding of Coa and Efb constructs were determined by comparing control wells with the ones where indicated amounts of mAbs were added. Average +- SEM of two independent experiments is represented.

**Supplementary Fig. 5 - Anti-CoaR antibodies do not inhibit Coa Fg binding**

Antibodies LIG40-A11 (left) and LIG40-D8 (right) were pre-incubated at the indicated amounts with the fixed amounts of Coa recombinant constructs (CoaF, CoaR0, CoaR at final concentration of 10nM, CoaC at 2nM) and then transferred on a Fg-coated ELISA plate. The remaining Fg-bound antigens were detected through their GST tag. Control wells in which no antibody was added were set to 100% and the residual binding of Coa constructs were determined by comparing control wells with the ones where indicated amounts of mAbs were added. Average +- SEM of two independent experiments is represented.

